# Single-cell sequencing of Entorhinal Cortex Reveals Wide-Spread Disruption of Neuropeptide Networks in Alzheimer’s Disease

**DOI:** 10.1101/2022.11.14.516160

**Authors:** Manci Li, Peter A. Larsen

**Author notes:** **Corresponding author:** Manci Li, BSc, Ph.D. candidate, Department of Veterinary and Biomedical Sciences, University of Minnesota, 1971 Commonwealth Avenue, St. Paul, MN 55108.

## Abstract

Alzheimer’s disease (AD) is a fatal neurodegenerative disease that involves early and significant neuropathological changes within the entorhinal cortex (EC). Many have reported on neuronal loss and synaptic dysfunction in the brains of AD patients and AD models. In parallel, abnormalities of neuropeptides (NPs) that play important roles in modulating neuronal activities are commonly observed in AD and other neurodegenerative diseases. However, the involvement of NPs has mostly been studied in the context of neurons; a cell type-specific examination of NP expression in AD brains is needed. Here, we aim to examine the NP networks in the EC of AD brains using single-nuclei and bulk transcriptomic data from other regions in the temporal cortex, focusing on the gene expression of NP and their cognate G-protein coupled receptors. We find that NP genes were expressed by all major cell types in the brain and there was a significant decrease in the quantity and the proportion of cells that express NPs in AD EC cells. On the contrary, the overall expression of GPCR genes showed an increase in AD cells, likely reflecting ongoing compensatory mechanisms in AD brains. In addition, we report that there was a disproportionate absence of cells expressing higher levels and greater diversity of NPs in AD brains. Finally, we established a negative correlation between age and the abundance of AD-associated NPs in the hippocampus, supporting that the disruption of the NP signaling network in the EC may contribute to the early pathogenesis of AD. In short, we report widespread disruption of the NP networks in AD brains at the single-cell level. In light of our results, we hypothesize that brain cells, especially neurons, that express high levels of NPs may exhibit selective vulnerability to AD. Moreover, it is likely AD brains undergo specific adaptive changes to fluctuating NP signaling, a process that can likely be targeted with therapeutic approaches aimed at stabilizing NP expression landscapes. Given that GPCRs are one of the most druggable targets for neurological diseases and disorders, we believe NP signaling pathways can be harnessed for future biomarkers and treatment strategies for AD.

## Introduction

Alzheimer’s disease (AD) is the most common neurodegenerative disease in people over 65, affecting over 50 million people worldwide and this number is expected to double every 20 years (1). Sporadic, late-onset AD makes up the majority of AD cases and it is associated with progressive cognitive, social, and functional decline after long prodromal cellular changes in the brain with the earliest and heaviest pathological changes occurring in the entorhinal cortex (EC) of AD patients (2). At the cellular level, major neuropathological characteristics of AD are neuronal and synaptic loss, glial activation, and amyloid plaques and neurofibrillary tangles formed respectively by misfolded Aβ and tau proteins (2). However, the pathological progression of AD is multifaceted, involving multiple cell types and mechanisms.

Neuropeptide (NP) signaling in the brain affects energy homeostasis, mood and motivation, sleep-wake states, immune response and inflammation, and ultimately behaviors (3,4). Despite decades of research, the exact definition of neuropeptides remains controversial (5–8). We use a broader concept of neuropeptide (NP) here adapted from Burbach (2011) (6); a neuropeptide is a small proteinaceous substance that can be but is not necessarily, synthesized by neurons and they are released through the regulated secretory route to influence cell signaling in neural tissues. NPs can exert their effects through autocrine, paracrine, and neurohormonal mechanisms. Many NPs have multiple defined cell surface receptors (9,10); most of these receptors are G protein-coupled receptors (GPCRs) found on the surface of cells and they transduce extracellular signals (e.g., hormones, odors, and neurotransmitters) into intracellular responses. GPCRs are among the most ‘druggable’ receptors due to their relatively easy accessibility and specificity in mediating downstream effects (11,12). Many drugs targeting GPCRs have been approved to treat neurological conditions and diseases and several GPCR-targeting drugs are in clinical trials for treating AD (11,12).

Due to the dynamic, pleiotropic, highly cell type-and region-specific, and often redundant nature of the NP signaling network, its breadth and specificity have been difficult to characterize (10). Recent studies leveraging single-cell sequencing (10) and micro-dissected bulk RNA-seq (8) have generated unparalleled physiological insights into the intracortical NP network. Additionally, advances in bioinformatics and machine learning (ML) have demonstrated great potential for discovering molecules and related gene expression signatures with diagnostic and therapeutic values for AD (13–18). While altered levels of components in NP signaling have been reported dispersedly in the EC of AD patients and various animal models (19), the extent of NP signaling disruption in EC has not been studied in depth at a single-cell level. To gain a better understanding of NPs and their roles in the pathophysiology of AD, a cell type-specific examination of NP expression and their cognate GPCRs (NP-GPCRs) in AD brains is needed.

In the present study, we conducted an in-depth transcriptome analysis focusing on NPs and their GPCR receptors in AD and control brains, using existing single-nucleus RNA-seq data from the EC and bulk transcriptomic data in nearby brain regions. We observed that at least one NP gene (*CCK*) was expressed by all major cell types investigated here (neurons, astrocytes, microglia, oligodendrocytes, and OPCs) in the EC of human brains. We report several NPs showing significantly lower transcript abundance and/or proportion of cells that express them in the EC of AD brains; they will be referred to collectively as AD-associated NPs (ADNPs). This is supported by bulk RNA-sequencing data in nearby brain regions of EC showing differentially decreased expression of many ADNPs and the predictive value of ADNPs demonstrated by ML. In contrast, the transcript abundance of NP-GPCRs increased in all five examined cell types. Further, we identified a disproportionate absence of cells expressing higher levels of NPs genes that also co-express more NP genes. Although most of these cells were GABAergic neurons, the expression of glutamate decarboxylases (GADs) was negatively correlated with the number of co-expressed NP in cells. Hypothesizing that the ADNPs contribute to the early pathogenesis of AD, we examined the correlation between the transcript abundance of ADNPs and age in the hippocampus and pancreas from the general population (bulk RNA-seq data from GTEx). We found a significant negative correlation between ADNP expression and age in the hippocampus but not the pancreas. In brief, we report widespread disruption of the NP network in AD brains at the single-cell level and our results support the hypothesis that NP network dysfunction participates in the early pathogenic processes in the development of AD.

Mitochondrial dysfunction and disrupted energy homeostasis at synapses occur early in AD (20,21). Synaptic transmission imposes high energy demands in neurons that can only be met by local mitochondria (21,22). Neurons shouldering more responsibilities in NP synthesis and excretion involve additional metabolic demands in macromolecule synthesis, transfer, and storage. This likely limits their ability to withstand temporary constraints in energy supply. In this context, our results deliver the perspective that GABAergic neurons expressing high levels and more types of NPs could be under high metabolic pressure and therefore selectively vulnerable to AD. We postulate that NP signaling pathways participate in early AD pathophysiology and thus can be harnessed for biomarker discovery and pharmaceutical development for AD as well as other neurodegenerative diseases.

## Material & Methods

### Obtaining and processing transcriptome data for AD analysis

#### Single-cell RNA-seq data

Raw data of single-cell RNA-seq generated from the entorhinal cortex of age- and sex-matched AD and control subjects (6 in each category, 12 total) was downloaded from a data repository (adsn.ddnetbio.com) provided by Grubman et al (23). Metadata is provided in detail in the original publication (23). Downstream analyses were performed using *Seurat* (v4.1.1) in RStudio (version: 2022.07.1, Build 554; R version: 4.2.1) (24,25). Cells were filtered based on the number of genes expressed and mitochondrial gene content; those with fewer than 200 and greater than 2500 detected genes as well as more than 5% of reads mapped to mitochondrial genes were filtered out. Normalization and scaling of data were performed as recommended by the *Seurat* pipeline. The count per million (CPM) matrix was generated using *NormalizeData* with the normalization method set to relative count (RC) and a scale factor of a million (1e6) as recommended by *Seurat*.

#### Bulk RNA-seq data

Raw count matrices from bulk RNA-seq data were downloaded from the Mount Sinai Brain Bank (MSBB) and Mayo RNAseq studies deposited in the ADKnowledge portal (26,27). RNA-seq data from Brodmann area 22 (BA22) and BA36 generated by the MSBB study was selected for this analysis due to their anatomical proximity to the entorhinal cortex. Criteria for AD in the MSBB study include CDR (Clinical Dementia Rating)>=1, Braak stage>=4, and CERAD (Consortium to Establish a Registry for Alzheimer’s Disease) score>=2. Individuals meeting the criteria of CDR<=0.5, Braak stage<=3, and CERAD score<=1 were considered controls. RNA-seq data from the temporal cortex produced by the Mayo RNAseq Study was used for machine learning as detailed below. Phenotype (AD or control) from the Mayo RNAseq Study was assigned in the original metadata. Samples recommended for removal were not included in this analysis. The final number of donors included for each condition from the above studies was summarized in Table 1. The denominated donor IDs from both studies were provided in Table S1. Protocols for RNA sequencing and data processing used to generate count matrices were documented in detail in the original studies (26,27).

**Table 1.** Individual samples included in the bulk RNA-seq analyses.

|  | Control | AD |
| --- | --- | --- |
| MSBB_BM22 | 42 | 114 |
| MSBB_BM36 | 45 | 98 |
| Mayo_TCX | 70 | 80 |

### Dimension Reduction

The top 2000 most variable features were identified during normalization. Both linear and nonlinear dimension reductions were then performed on scaled data by Principal Component Analysis (PCA) of variable features and Uniform Manifold Approximation and Projection (UMAP) analysis, respectively. The first nine dimensions were used for UMAP analysis as determined by the inflection point on the elbow plot (Fig S1). Resolution for the *FindNeighbors* function was set at 0.2 and the *FindClusters* function resulted in 10 clusters that were in good agreement with the association scoring based on the *BRETIGEA* brain cell markers (detailed below; Fig S2).

### Cell identification

Methodology for identifying cell types was adopted and modified from Grubman et al (23). In short, *AddModuleScore* in *Seurat* was first used to calculate association scores using lists of brain cell type-specific markers from the *BRETIGEA* package (v1.0.3) (24,28). Six major cell types were identified using 50 markers for each cell type as included in *BRETIGEA*: astrocyte (ast), microglia (mic), neuron (neu), oligodendrocyte (oli), oligodendrocyte precursor cells (opc), and endothelial cells (end). Cells were labeled based on the highest association score except 1) when they were considered unidentified (lowest 5% in each cell type) or 2) when they were considered hybrid (the highest and the second highest score were within 20% of each other. In addition, we labeled cell clusters generated from *Seurat* based on the *BRETIGEA* scoring (Fig S2). Cell identification based on clustering was adopted (Fig S3A) but unidentified and hybrid cell labels were kept from the first set of identification. Additionally, cells in cluster 8 were labeled as unidentified as they had low association scores for both neurons and oligodendrocytes. A summary of cell identification for each sample is provided in Table S2. Successful cell type identification was confirmed by a heatmap of differential gene expression of cell type markers among cell groups (Fig 1C). Endothelial, hybrid, and unidentified cells were excluded from the downstream analysis related to NPs and their receptors.

**Fig 1.**
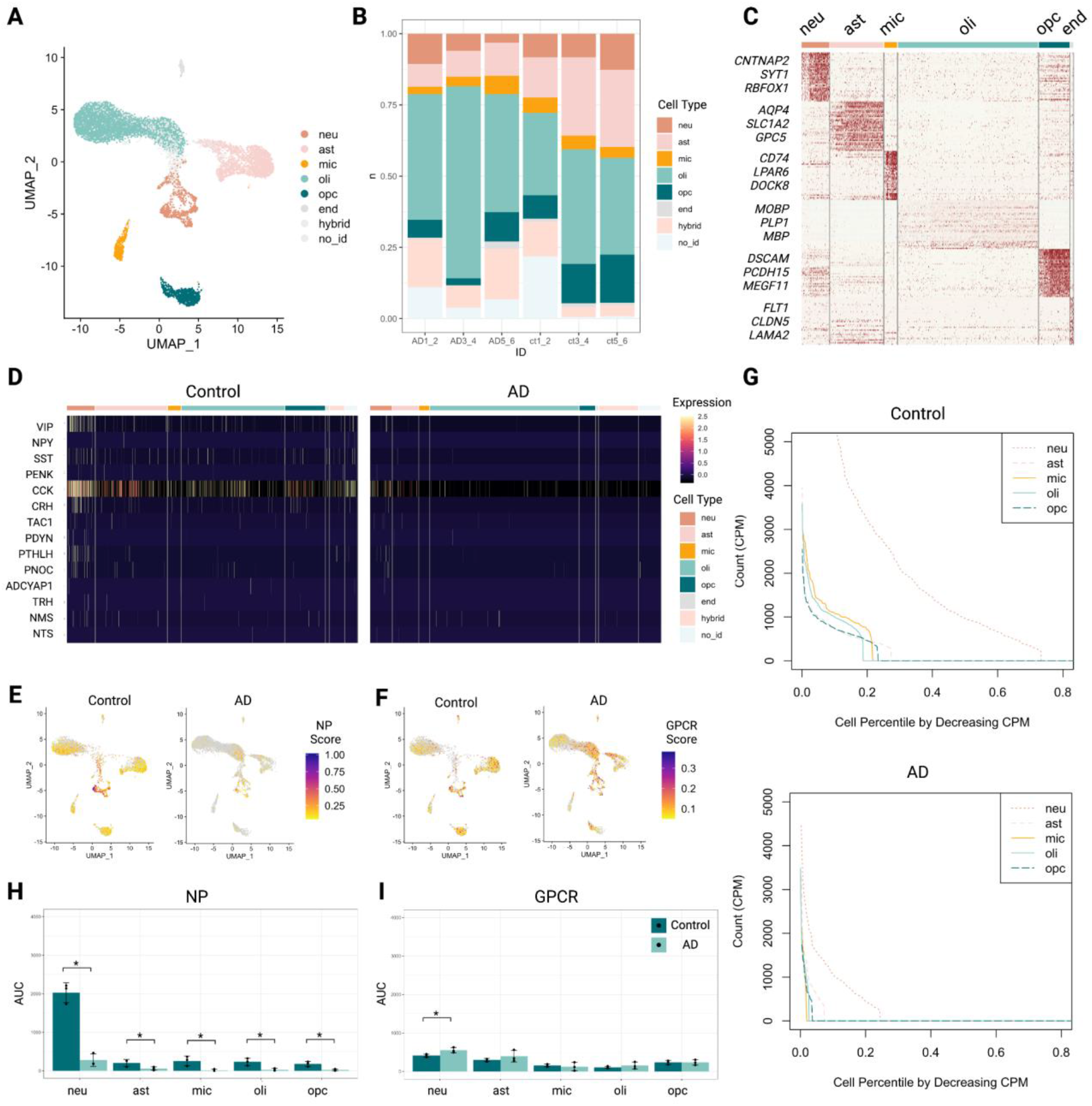
Decreased expression of neuropeptides (NP) revealed by single-cell sequencing data. (A) Cell identity shown on the dimension reduction plot. (B) Cell composition of grouped replicates (two biological replicates per group). (C) Differential gene expression confirmed the identification of cell marker genes for each assigned cell type. (D) Heatmap showing expression levels of selected NP genes in each cell type in control and AD brains. (E) Gene module score for selected NPs calculated from control brains visualized on the dimension reduction plot. (F) Gene module score for selected NPs calculated from AD brains visualized on the dimension reduction plot. (G) The total relative count of all selected NPs plotted against the cell percentile ranked by decreasing cell count in selected cell types in control and AD. (H) The area under the curve (AUC) of NP transcription for selected cell types in each grouped replicate in control and AD. One-tailed Wilcoxon signed-rank test was used. (I) The AUC of GPCR transcription for selected cell types in each grouped replicate in control and AD. One-tailed Wilcoxon signed-rank test was used. *, p<0.05; no_id, cell assigned as no id; neu, neuron; ast, astrocyte; mic, microglia; oli, oligodendrocyte; opc, oligodendrocyte progenitor cells; CPM, count per million; GPCR, G-protein coupled receptors.

### Differential gene expression

#### Single-cell sequencing data

Differential gene expression (DGE) was first used to provide unbiased verification of results from cell identification. The top marker genes in each putative cell type were determined by the *FindAllMarkers* function; the Wilcoxon rank-sum test was used, and the False Discovery Rate (FDR)-adjusted p-value (p-adj) was set to be less than 0.01. The top 35 marker genes ranked by the average 2-fold difference from each putative cell type were used as input to validate cell identification visualized by *DoHeatmap*. Cell markers for validation used by Grubman *et al* were successfully reproduced (23). The same function and settings were applied to obtain cell-specific differentially expressed genes between AD and control subjects.

#### Bulk RNA-seq data

Raw read counts from BA22 and BA36 generated by the MSBB study were selected for DGE analysis using the DESeq2 package (v1.30.1) in RStudio (25,29). For multiple comparison correction, Benjamini-Hochberg correction was applied to calculate FDR-adjusted p-values (p-adj). The significant term was determined by using a cut-off of 0.05 (p-adj<0.05). Lists of significantly decreased genes were saved and used as input for STRING analysis later.

### Scoring of NP and receptor transcription

Lists of NP and their receptors were manually searched and integrated from several peer-reviewed sources (10,30–33). Only NPs with over 9200 CPM across all cells were included in the downstream analysis. *AddModuleScore* in *Seurat* was used to calculate association scores using gene lists of NP and their receptors provided in the supplementary material (Table S3). Results were visualized by *FeaturePlot_scCustom* in the *scCustomize* package with the cut-off score set at 0.05 (34).

### Analyzing the rate of transcription/synthesis

Collective CPM of included NP or receptors for each cell in descending rank order was plotted along an increasing cell population percentile for each cell type. The area under the curve (AUC) was calculated using *trapz* function in the *pracma* package (v2.3.8) for each phenotype and each sample (35). As reasoned by Smith et al., the abundance of transcripts for NP and their receptors can be construed as the potential for the rate of synthesis of the encoded protein (10). Here, AUC formed by CPM vs cell percentile can, therefore, be used as the proxy for the relative rate of target transcription at least if not peptide and protein synthesis.

### STRING analysis

Genes with significantly decreased expression in the BA22 and BA36 of AD brains were selected and uploaded to STRING (Search Tool for the Retrieval of Interacting Genes/Proteins) (v11) (36). STRING was used to construct functional protein association networks for probable gene products; Euclidean distances were used to cluster gene products. Local network clusters were downloaded from STRING analysis.

### Machine learning

Machine learning (ML) was carried out using the *caret* package (Classification And REgression Training, v6.0-92) in RStudio using temporal cortex transcriptome generated by the Mayo RNASeq Study (25,27,37). Donors were split into training and testing cohorts before ML model training; there was no overlap between donors in the two cohorts. Centering and scaling were done by *preProc* in *caret*. ML algorithms selected here included *‘glmnet’* (General Linear Model fit via penalized maximum likelihood, GLMNET), *‘ranger*’ (Random Forest, RF), and ‘*svmRadial*’ (Support Vector Machines with Radial Basis Function Kernel, radial kernel SVM). Cross-validation was set as 10 for all controls. GLMNET fits generalized linear and similar models via penalized maximum likelihood (38). Two hyperparameters were tuned: alpha (mixing percentage) and lambda (regularization parameter) were customized using *expand.grid*. Alpha was optimized between 0 and 1. A hundred values of lambda between 0.0001 and 1 were fit per alpha on the training data. For the final mode using ADNP as input features, alpha was set at 0 and lambda at 0.0405. RF uses an ensemble of classification trees to classify. The hyperparameter *mtry* (number of randomly selected predictors) required by RF was tuned by default. *Mtry* was optimized to 19 for the ADNP RF model. Radial kernel SVM generates non-linear decision boundaries to separate non-linear data. Two hyperparameters, Cost (C) (the penalty for misclassification) and sigma (influence of one training example), were optimized with a tuning length of 15. The final SVM model using ADNPs had a sigma of 0.0199 and C of 0.5. Code for ML application is detailed in the reference manual of the *caret* package (37). ML models were trained using the training cohort and then applied to the testing cohort to obtain the risk score for each sample in the testing cohort. The risk scores were then compared to the true disease status of the samples to evaluate the accuracy and generate ROC curves. Lastly, feature selection was performed to extract the top 15 most valuable ADNPs for each model. *VennDiagram* package (v1.7.3) was used to visualize the consensus among models (39).

### Transcriptome data and analysis for studying ADNP changes in the general aging population

RNA-seq data matrices of the human hippocampus and pancreas from the general population containing normalized transcript count, transcript per million (TPM), were downloaded from the GTEx portal (40). Details of data generation and processing were described in the publication by the GTEx Consortium (41). Individuals without complete metadata (age, sex, death classification) were excluded from our analysis. In addition, those that scored 3 and 4 (intermediate and slow death) on the Hardy Score for death classification were excluded. Since it is well known that brain development continues throughout the early 20s (42), subjects in the age bracket of 20-29 were also excluded. Further, individuals that showed no expression of one or more NPs were excluded. Due to the tissue specificity of NPs, 8 ADNP genes (*PNOC, NPPC, TAC3, NXPH1, PDYN, NXPH2, CBLN2, HCRT*) not expressed at the RNA level in the pancreas were not used in the ADNP analysis for the pancreas. Ultimately, 38 ADNPs from 74 individuals (hippocampus) and 30 ADNPs from 141 subjects (pancreas) were included in this analysis aiming to identify the association between age and NP expression using bulk RNA-seq data. All normalized transcript counts were log2 transformed. Pearson correlation test was performed whenever correlation was tested; the significance cut-off was set at 0.05 for the sum of transcript count and 0.1 for individual ADNP transcript. One-tailed Wilcoxon signed-rank test was used to test the difference in the correlation coefficients derived from the sum of three random genes from the ADNP pool (38 for hippocampus, 30 for pancreas) and five thousand from the non-ADNP transcript pool (proportional to ~3:38); significance cut-off was set at 0.05.

### Additional Statistical Information

The percentile or proportion of cells is based on one specific cell type in two biological replicates (2 individual/count matrices), unless otherwise stated. For example, the proportion of VIP-expressing neurons in AD1_2 was calculated as (neuron count in AD1_2 expressing VIP)/(neuron count in AD1_2). Wilcoxon’s rank-sum test was conducted as one- or two-tailed versions as stated in the text based on different biological questions and hypotheses. Significant terms were set at 0.05 or 0.1 depending on biological questions as specified in the text.

## Results

### Decreased expression of neuropeptides (NP) in AD entorhinal cortex

To investigate the expression of major NP and their cognate receptors in different cell types in the AD EC cortex, we reanalyzed single nuclei sequencing data generated by Grubman et al (23). The dataset consisted of six AD patients and six sex- and age-matched controls; these individuals were divided into groups of 2 for each condition (e.g., AD1_2, AD3_4, AD5_6, control1_2, control3_4, control5_6). Details of the metadata, including the *APOE* genotype, can be found in the source publication (23). We conducted independent quality filtering, dimension reduction, and cell identification. After quality filtering (Methods), we included 14279 cells (AD: 7333; control: 6946) and the median detected gene per cell was 689. UMAP was used to visualize the transcriptome and assist in cell identification (Fig 1A). Cells from AD patients were well-separated from those in controls (Fig S3B&C). In addition to unsupervised classification (UMAP), previously established cell-type-specific gene sets (included in the BRETIGEA package) were also used to identify cells (28). Nuclei were separated into several clusters mapped to the six major cell types in the brain as established by BRETIGEA, including microglia, astrocytes, neurons, oligodendrocyte progenitor cells (OPCs), oligodendrocytes, and endothelial cells (Fig 1A). We also adopted the ‘hybrid’ and ‘unidentified’ cell types as described by Grubman et al with modifications. Specifically, cells were labeled as ‘hybrid’ if the difference between the two highest cell type scores were less than 20%, or as ‘unidentified’ if the highest cell type score was within the lowest 5% (detailed in Method). In addition, we labeled cluster 8 from UMAP as ‘unidentified’ as it had low scores for both neurons and oligodendrocytes (Fig S3A). Since we would not include hybrid, unidentified, and endothelial cells for downstream analysis, implementation of stricter inclusion criteria would not impact further investigations. The proportions of cell type between control and AD phenotypes were similar, except for the marginally higher proportion of hybrid cells in AD brains (p=0.081) (Fig 1B&S3D). To confirm the validity of our cell identification, we used differential expression between cell types and reproduced marker genes reported by Grubman et al, specifically *CNTNAP2, SYT1*, and *RBFOX1* for neurons*, AQP4, SLC1A2*, and *GPC5* for astrocytes, *CD74, LPAR6*, and *DOCK8* for microglia, *MOBP, PLP1, MBP* for oligodendrocytes, *DSCAM, PCDH15*, and *MEGF11* for OPCs, as well as *FLT1, CLDN5*, and *LAMA2* for endothelial cells (Fig 1C).

Several NPs (e.g., CCK, neurotensin, PACAP, neuropeptide Y) showed decreased gene expression and levels in multiple tissue types of AD patients and animal models, such as blood and brain tissues (19,43). Therefore, we predicted that there would be a decreased expression of several NPs in the EC of AD patients at the single-cell level as well. We additionally investigated the NP-GPCRs as NPs can exert functions through autocrine and paracrine signaling, examining the change of expression of these GPCRs may reveal local adaptive changes of brain cells to NP loss. We first examined and visualized the expression of *NP* and their cognate receptors selected by Smith et al. In the control EC, all cell types expressed multiple NPs, and neurons have the highest expression of selected NPs (Fig 1D). Interestingly, *CCK* is highly expressed by all five cell types (Fig 1D). As predicted, the expression selected NP is much sparser in AD brains; however, the extent of the difference observed in AD brains was surprising (Fig 1D). Differential gene expression between AD and control among different cell types showed that the expression of several NPs (*CCK, VIP, CRH, SST*, and *PTHLH*) was significantly decreased in neurons, and *CCK* was significantly decreased in all included cell types (Table S3; Fig S3E). *CCK* encodes for a neuropeptide precursor that is processed to generate several neuropeptides, for example, cholecystokinin (CCK) −8, −12, and −33 in the gastrin/cholecystokinin family of proteins. Besides its gastrointestinal functions, CCK has important roles in the CNS, such as feeding behavior, mood regulation, as well as learning and memory processes (44–46). CCK is decreased in the brain and CSF of AD patients; the latter is also associated with tau and phosphorylated tau in the CSF. Similar to CCK, neuropeptides including VIP (47), CRH (48,49), and SST (50) were all documented to be altered in AD and could exert neuroprotective effects against AD (19,49,51–55). These NPs have major functions related to abnormalities observed in AD, such as regulating mitochondrial morphology and/or activity (56–60), participating in circadian processes (52,61–67), and reducing neuroinflammation and/or glial activation (53,68–72). PTHLH is known to regulate the development of the skeletomuscular system (73), but its neurological functions are less known although it is expressed throughout the brain (proteinatlas.org; accessed 2022 Sep). It has not been studied extensively for AD, except a recent report identifying differential methylation in males and females with AD (74). In contrast to the drastic difference observed for NP expression between AD and control subjects; none of the listed GPCRs were differentially expressed (Table S3).

To visualize the overall expression of NP and associated GPCRs in control and AD cells, we used the selected genes of NPs and GPCRs as input features to calculate module scores in the *Seurat* package, which compares the average expression level of input features to randomly selected genes. Results show decreased *NP* expression in all cell types, especially neurons, in AD and there was no apparent change in GPCR expressions (Fig 1E&F). To quantify the difference that we observed in module scores and derive a statistically supported conclusion, we plotted the relative count of transcripts (count per million, CPM) against the cell percentile ranked in decreasing order for each cell type, including neurons, astrocytes, microglia, oligodendrocytes, and OPCs. In this way, the area under the curve (AUC) in the plot can be used as a proxy for the relative rate (capacity) of NP and GPCR transcription in each cell. In control cells, ~80% of neurons and ~20% of the rest of the cell types transcribe NP express; these numbers dropped sharply to ~20% and ~5% in AD cells, respectively (Fig 1G). We calculated and compared the AUC for each grouped biological replicate (containing 2 subjects) for control and AD subjects and we found that the overall expression of the listed NPs was significantly decreased in the five cell types included here (Fig 1H). However, this was not the case for GPCRs; in fact, the expression of GPCRs in AD neurons was significantly higher than in controls (Fig 1I). This analysis provided quantitative support to the observation that there is a decreased expression of NPs but not GPCRs in AD EC cells.

### Decreased NP cell population in AD entorhinal cortex

To investigate NP changes further, we decided to include other NPs and their GPCR receptors in our analysis. Since the concept of NPs— endogenous peptides affecting and interacting with the nervous system through binding to cell surface receptors that can be but are not necessarily synthesized by neurons—is in part specific to this study, we used several peer-reviewed sources (10,30–33) to generate the gene lists for NP and their GPCRs, and only NP genes expressed with a total CPM over 9200 (the lowest total CPM examined in those reported by Smith et al.) and their cognate GPCRs were included for analysis. We also included neurexophilins, a family of neuropeptide-like glycoproteins, in the list. As a result, 49 NP genes and 50 GPCRs were selected (Table S4).

Next, we repeated the module visualization, cell-type specific differential gene expression, and AUC calculation for the more comprehensive gene lists. Again, we observed a lower expression of NPs, especially in neurons and OPCs, but not GPCRs in AD cells (Fig 2A). We also identified additional differentially expressed genes on the NP list, including *SCG2, IGF1, SCG5, NXPH1, CHGA, CHGB*, and *SCG3* in neurons, *SCG3* in astrocytes, and *NXPH1* in OPCs (Fig 2B&S4A). *SCG2, SCG3, SCG5, CHGA*, and *CHGB* encode for neuropeptides precursors/neuropeptides secretogranin II, Secretogranin III, 7B2, chromogranin A (CGA), and chromogranin B (CGB) that will be proteolytically processed into neuropeptides in the chromogranin–secretogranin family (75). These peptides are major constituents of and often markers of dense core vesicles (DCV) that store neuropeptides (76). Our finding signifies that the disruption of NP signaling in AD is likely widespread (e.g., not limited to a couple of NPs). In contrast to the changes in NP expression, differential expression analysis of GPCRs showed limited alterations only in astrocytes and OPCs in AD EC (Table S5).

**Fig 2.**
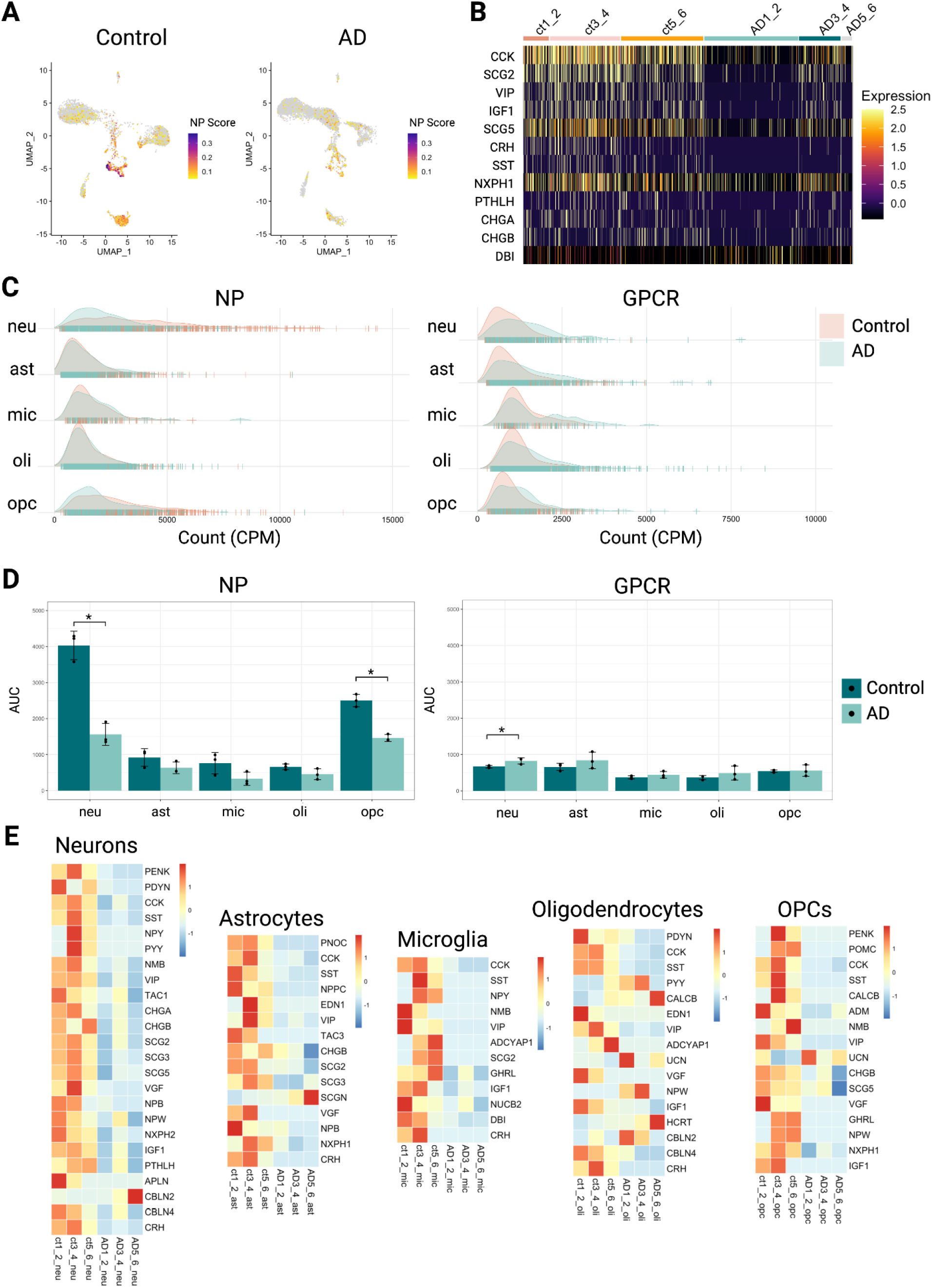
The population of NP-expressing cells was decreased in the AD entorhinal cortex. (A) Gene module score calculated from control and AD brains using genes in the expanded NP list was visualized on the dimension reduction plot. (B) Heatmap showing expression levels of differentially expressed NP genes in neurons from control and AD brains. (C) Distribution of control and AD cells in each cell type by relative counts of total NP and GPCR transcripts. NP, neuropeptides; GPCR, G-protein coupled receptors. (D) The area under the curve (AUC) of NP and GPCR transcription compared for selected cell types in each grouped replicate in control and AD. One-tailed Wilcoxon signed-rank test was used. *, p<0.05; NP, neuropeptides; GPCR, G-protein coupled receptors. (E) Cell type-specific heatmaps showing the NP genes expressed by significantly changed (One-tailed Wilcoxon signed-rank test, p<0.1) proportions of cells for each grouped biological replicate. Genes expressed only in one grouped replicate were excluded. Relative proportion = number of cells expressing gene target for each cell type per biological replicate/number of cells in the cell type per biological replicate.

Significantly decreased expression of the NPs in AD EC cells reported here agrees with decades of research surrounding synaptic pathology in AD (77–79); secretoneurin, CGA, and CGB are colocalized with plaques and have been shown to decrease in the EC of AD brains (78). Our findings here likely reflected secretory vesicle dysfunction and/or a loss of neurites and neurons, perhaps especially in cells that store and secrete large quantities of neuropeptides, in the EC of AD subjects. *IGF1* encodes for insulin-like growth factor 1 (IGF-1)— a highly pleiotropic peptide synthesized by both neurons and glial cells that promotes neurogenesis and regeneration (80–82), modulates synaptic metabolism and function (83–85), protects against neuroinflammation through regulating astrocytes and microglia activities, and potentially decreases blood-brain barrier permeability in the CNS (86). Higher levels of IGF-1 have been associated with decreased risk of AD and IGF-1 is significantly decreased in the serum of AD patients (87). The drop in IGF-1 level has been proposed to mediate neuronal insulin resistance and the development of hallmark neuropathological changes in AD (88–91). Neurexophilin-1, encoded by *NXPH1*, is localized to presynaptic terminals in neurons and forms a tight complex with α-neurexins (92), a transmembrane cell adhesion protein, to modulate short-term plasticity and the distribution of GABA receptors in selective subpopulations of synapses (93). An SNP (rs6463843) flanking *NXPH1* was identified to be associated with global and regional grey matter density loss in AD patients in a GWAS of neuroimaging phenotypes (94). In summary, the differential expression of NPs identified here is in congruence with existing literature on AD describing synaptic and metabolic dysfunction in neurons, especially impaired synaptic transmission mediated by DCV.

As we examined the distribution of total NP and GPCR for each cell, we found that more neurons and OPCs in control cells were expressing a higher level of NP with more variance while GPCR expression in all five AD cell types seemed to have also increased in both the mean and variance (Fig 2C). This is an interesting observation especially if one considers the increased proportion of hybrid cells in AD (23); the abnormality in the distribution then seems to suggest that there was a decrease in subtype diversity and transcription capacity in neurons and OPCs in terms of NP synthesis while more cells were adapting to express higher levels of GPCRs perhaps as a compensative strategy to the loss of their NP ligands in AD. Compared to the smaller list analyzed previously, OPCs in AD EC, in addition to neurons, also expressed NP at a significantly lowered rate in AD cells when compared to controls (Fig 2D&S4B). GPCRs showed a marginally opposite change to NP, where most cell types in AD EC expressed GPCRs at a slightly but significantly higher rate (Fig 2D).

Because the method used here for differential gene expression analysis more accurately reflects the average gene expression per cell rather than altered cell proportions and therefore cannot fully represent the abnormalities of NP expression in AD, we set out to identify NPs and GPCRs expressed by altered proportions of the cell population. We calculated the relative proportions of cells that express individual gene targets of interest for each grouped biological replicate (number of cells expressing gene target for each cell type per biological replicate/number of cells in the cell type per biological replicate) and plotted the ones that showed a significant decrease in AD as cell type-specific heatmaps. As expected, this analysis revealed that more NPs were expressed by abnormally lower levels of cells in AD (Fig 2E). Besides oligodendrocytes, all four cell types (neurons, astrocytes, microglia, and OPCs) showed a near-consistent decrease in the NP-expressing populations in AD (Fig 2E). Noticeably, the decrease in cell populations expressing those genes previously highlighted in this analysis (*SST, VIP, CRH, NPY, IGF-1, NXPH1*, and genes encoding for neuropeptides in the chromogranin-secretogranin family) was no longer limited to neurons, supporting the idea that AD EC has lower capacity in synthesizing and secreting NPs. Additionally, the population of cells expressing *VGF* and members of the endogenous opioid precursors (*PENK, PDYN, PNOC, POMC*) was decreased in four out of five cell types. *VGF* encodes for neuropeptide precursor VGF, which is also a member of the secretogranin family that functions in DCV and can be proteolytically processed into at least 12 VGF-derived peptides (95–97). Decreased VGF level and its dysregulation in AD is well-established and recently reviewed (98). Endogenous opioids have broad effects on the brain, including regulating sensory, emotional, motivational, and cognitive functions, as well as promoting neuroinflammation (99,100). While the role of the endogenous opioid system in AD pathogenesis remains unclear, our results suggest that endogenous opioids may participate in the development of AD by coordinating the communication among multiple brain cell types. While we showed previously that the overall transcript abundance increased for GPCRs in neurons and OPCs, cell populations expressing GPCRs consistently decrease in these cell types in AD, and far fewer GPCR genes (six in neurons and four in OPCs) were abnormally expressed (Fig 2E&S4C). Because the abundance of these GPCR genes did not significantly differ between control and AD EC, this observation complements our previous finding from examining the distribution of transcript abundance where there was an increased cell population expressed more GPCRs in response to the decreased NP signaling (Fig 2C&D); some cells were likely expressing more of the identified GPCRs to the loss of GPCR expression by other cell populations.

### Dysregulation of NP network in AD supported by bulk RNA-seq datasets

Because the chosen single-cell dataset consisted of limited biological replicates (6 individuals per condition) and sequencing depth, we wanted to confirm our finding of widespread NP expression decline with other RNA-seq data obtained near EC that comprised more subjects before conducting further analysis. To this end, we chose the bulk RNA-seq data generated from postmortem brain tissue collected through the Mount Sinai VA Medical Center Brain Bank deposited on the AD Knowledge Portal. Brodmann area (BA) 22 and 36 were selected due to their proximity to EC (Fig 3A). We identified the genes with significantly decreased expression in AD compared to controls and used them as input for STRING, which is a database of known and predicted protein-protein interactions, including direct (physical) and indirect (functional) associations (36). Agreeing with our results from single-nuclei sequencing, many NP genes but not their cognate GPCRs were expressed at a significantly lower level in AD brains shown by bulk RNA-seq in both regions near EC in the temporal cortex. In addition, protein products of genes with significantly decreased expression formed association clusters surrounding components of neuroactive ligand-receptor interaction and neuropeptides (Fig 3B). This gave us confidence that although the single-cell data analyzed here was generated from limited subjects, the data should reflect the key changes regarding NP expression in AD.

**Fig 3.**
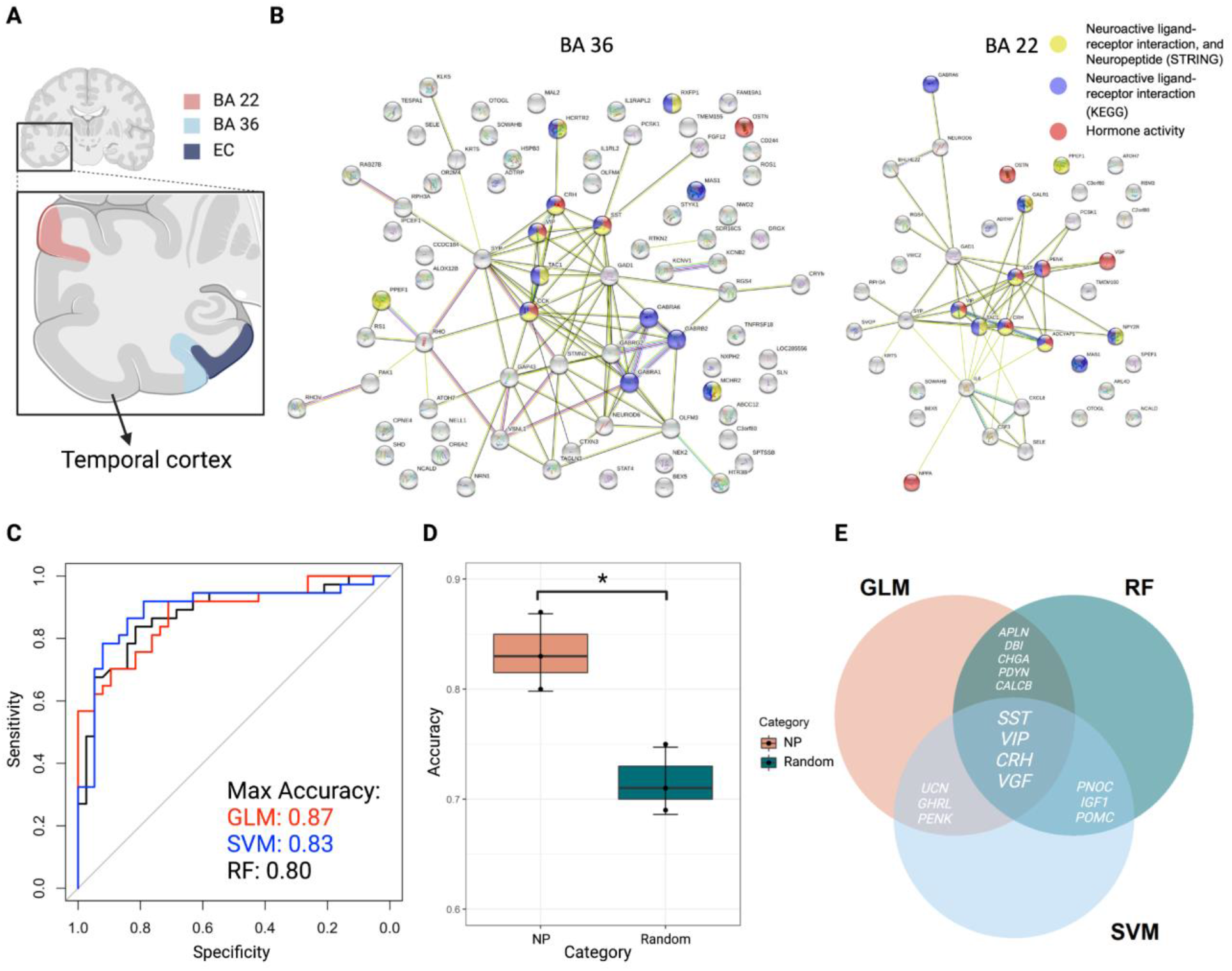
Analyses of bulk RNA-seq data from the temporal cortex highlight importance of NPs in the molecular pathology of AD. (A) The neuroanatomical diagram of the brain tissues used to generate RNA-seq data included in this study. (B) STRING analysis showing gene product interaction networks formed by significantly decreased genes in AD from Brodmann Area 36 (BA36) and BA22. RNA-seq data from the MSBB (Mount Sinai Brain Bank) study was used. (C) ROC (Receiver Operating Characteristic) curves of three different machine learning (ML) models—GLM, RF, and SVM—using ADNPs as input variables applied to bulk RNA-seq data generated from the temporal cortex. Data collection was performed by the Mayo RNASeq study. (D) Comparing the maximal accuracy of ML models generated using ADNPs vs the same number of random non-ADNP genes as input features. One-tailed Wilcoxon signed-rank test was used. *, p=0.05. (E) Venn-Diagram of ML-driving ADNPs extracted from three different ML algorithms. GLM, glmnet; RF, random forest; SVM, support vector machine.

As a decreased expression of NPs can be observed in EC, BA36, and even BA22, we then speculated that the disruption of the NP network could be more ubiquitous and reflected at the temporal cortex level for AD patients. To show this, we applied common machine learning models (random forest (RF), logistic regression, and supportive vector machines (SVM)) used on transcriptome (13,101–104) to bulk RNA-seq data generated from the temporal cortex (TCX) (27). The list of ADNPs was generated by combining all the ADNPs discovered to have altered expression patterns at the cell population level from different cell types shown in Fig 2E. We found that all three models could use the ADNPs to predict AD vs C with an accuracy of over 80% (Fig 3C) and this is significantly higher than the accuracy of ML results generated by random genes expressed in the TCX (Fig 3D). NP signaling can be redundant, therefore, to identify the most valuable NP expression that could distinguish AD and control, we extracted feature importance–which represents how useful input genes are at predicting AD status–from all three ML models and selected the top 15 most useful genes identified by each model. Agreeing with our findings from the single-cell and the MSBB bulk RNA-seq data, *SST, VIP, CRH*, and *VGF* were considered valuable for prediction by all three models (Fig 3E). All members of the opioid family (*PENK, POMC, PDYN, PNOC*) were considered useful in two models (Fig 3E). *CHGA*, a marker gene for DCV, was also highlighted by two models (Fig 3E). In sum, we used two independent sets of transcriptome data to support our findings from the in-depth analysis of the NP signaling network using single-nuclei RNA-seq data. We conclude that results from our single-cell sequencing analysis are reflected at the bulk RNA-seq level and the disruption of the NP signaling network in the AD temporal cortex is widespread.

### Disproportionate absence of high NP expression in cells from AD entorhinal cortex

We observed more consistent and widespread changes in NP expressions in comparison to GPCRs, therefore we decided to focus on investigating the expression of NP patterns at the single-cell level, especially the rate/quantity and types of NP expressed per cell in the EC of AD and control subjects. We believed that this line of investigation could shed light on the contributing factors underlying the significant decrease in NP expression in AD cells.

Since we noticed that lowered expression of *GAD1* was tightly associated with the expression of NP in bulk RNA-seq, we confirmed that *GAD1* was also expressed at a significantly lower level in AD neurons from the single-nuclei data focused on here (Table S6). This is not surprising as the level of GABA is known to be decreased in the temporal cortex of AD patients (105,106), indicating a dysfunctional GABA system in AD. GABAergic neurons, also known as GABAergic interneurons, often synthesize and secrete multiple types of neuropeptides (107). Considering that cells with higher transcription activity and storage of neuropeptides might be facing greater metabolic demands and our finding that there was a decrease in cell populations that express higher levels of NP (Fig 2C) in AD, we wondered if some GABAergic cells in the EC shouldering more responsibility to produce NP were therefore disproportionately dysfunctional/lost in AD. We also questioned if these cells simply dedicated more transcripts to producing a higher proportion of specialized NPs or if they also co-expressed multiple NPs.

To address these questions and provide some insight into the NP cells that are lost in AD, we selected cells that are producing over 7000 CPM of NP, which accounts for ~0.7% of the total transcripts produced by the cell. These cells will be referred to as high NP-producing (HNP) cells hereon. We found that there were only 14 HNP cells in this AD EC while control brains had 107 (Fig 4A; Table S7). As expected, there was a disproportionately higher absence (p<0.05) of HNP cells, which were mostly neurons, in AD (Fig 4B; Table S8). Next, we examined the NP co-expression pattern in control and AD EC cells. In control cells, all five cell types showed evident NP co-expression; neurons and OPCs formed right-skewed bell curve distribution while the proportions of NP co-expression decreased as the number of co-expressed NPs increased in other cell types (Fig 4C). In contrast, the distribution of AD neurons based on the number of co-expressed NPs lost the bell curve shape completely, with ~1/3 of the cells not expressing any studied NPs, suggesting that many AD neurons have decreased ability to co-express NPs (Fig 4D). Complementary to the previous analysis where we identified NPs expressed by an altered population of cells, the proportions of cells co-expressing NPs also changed. Specifically, we found a significant increase in the proportions of cells that don’t produce any analyzed NP in all five cell types from AD (Fig 4E).

**Fig 4.**
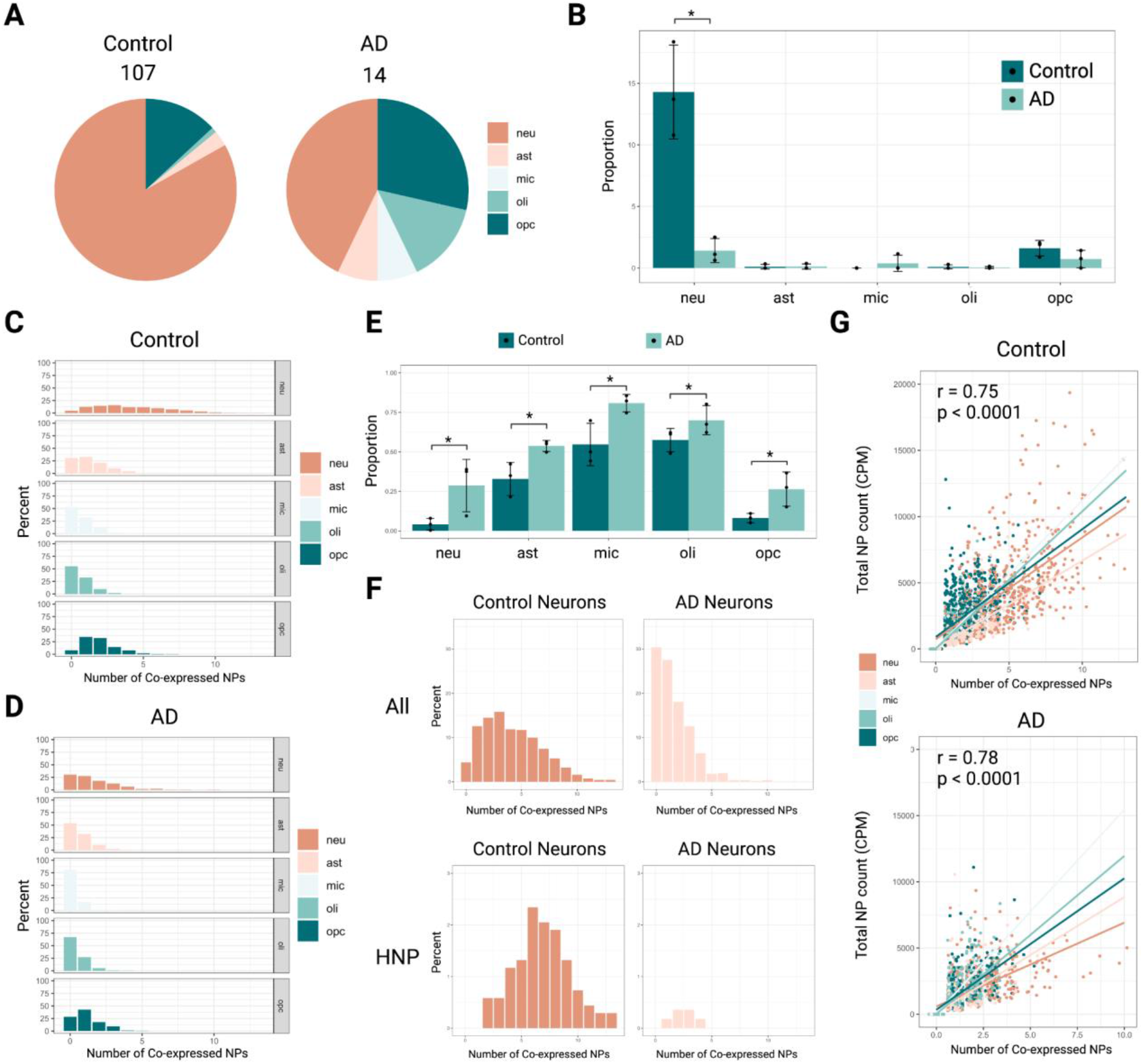
Analyses of high NP-producing (HNP) cells and NP co-expression highlight a disproportionate absence of HNP cells in the AD entorhinal cortex (EC). (A) Pie charts demonstrating the quantity and proportion of HNP cells in control and AD EC. (B) Cell type-specific comparison of the proportion of HNP cells in control and AD EC. One-tailed Wilcoxon signed-rank test was used. *, p<0.05. (C) Cell type-specific distribution of control EC cells based on the number of co-expressed NPs. (D) Cell type-specific distribution of AD EC cells based on the number of co-expressed NPs. (E) Cell type-specific comparison of the proportions of cells that express no detectable NPs in control and AD EC. One-tailed Wilcoxon signed-rank test was used. *, p<0.05. (F) Distribution of all neurons and HNP neurons based on the number of co-expressed NPs in control and AD. (G) Significant correlation between the sum of the relative number of NP transcripts and the number of co-expressed NPs shown in both control and AD EC cells. Pearson’s test of correlation was used. r, correlation coefficient.

As most HNP cells were neurons, we focused on the expression pattern of these cells. Surprisingly, we found that HNP neurons were also co-expressing more genes encoding for NPs compared to the general population of neurons (Fig 4F). Although there were very few neurons producing NPs at a comparable level in AD (Fig 4F), these cells also produced at least one NP included in this study (Fig 4F). Based on these observations, we hypothesized that cells co-expressing more types of NPs were also dedicating higher proportions of their transcriptional capacity to NP expression. To test this idea, we calculated the correlation between the total CPM of NP transcripts and the number of co-expressed NPs. Despite the scarcity of HNP cells in AD, we discovered that there was a strong and significant positive correlation between the relative transcription capacity dedicated to NP and co-expression in AD (p<0.0001; r=0.75) similar to controls (p<0.0001; r=0.78) (Fig 4G), suggesting that co-expression of NPs likely requires the higher dedication of transcription capacity, and this requirement is still being fulfilled in existing HNP AD cells.

The brain has long been known to be energetically expensive, consuming 20% of the body’s energy while measuring only ~2 to 2.5% of the weight with no capacity to store glucose locally. Although some of this energy consumption is due to brain electrical activity, most are devoted to basal metabolism (108). A recent study showed that ATPases (V-ATPases) compensating for a resting H+ efflux from the synaptic vesicle (SV) lumen contributed significantly to basal metabolic processes in glutaminergic neurons, highlighting one potential source of the metabolic vulnerability due to high energy consumption (108). While basal energy consumption has not been measured for SVs in GABAergic neurons or dense core vesicles (DCV) containing NPs, it is known that V-ATPases are responsible for initiating the acidification of GABA SV (109) and are active on DCVs (110). Because NP-expressing neurons are often GABAergic, our results could lead to the hypothesis that GABAergic neurons, especially those producing a large number of NPs, have higher metabolic demands to generate, maintain, and secrete numerous NPs in addition to GABA, therefore are selectively vulnerable to metabolic stress. However, since *GAD1* also had significantly lower expression in AD neurons (p<0.0001) and storing neurotransmitters was expected to be energetically costly (108), aspects of GABA processing, storage, or activity rates may be driving the basal energy demands of neurons rather than the quantity and types of NPs that are produced despite our analysis surrounding NP expression in AD cells. Therefore, we subsequently investigated whether there is a correlation between *GAD1* and *GAD2* (genes encoding for GADs) expression and the disproportional absence of neurons co-expressing multiple NPs. If metabolic demands behind synthesizing and storing GABA is the key feature driving the absence of NP-expressing cells in EC of AD subjects, we would expect a significantly positive correlation between *GAD1* and/or *GAD2* expression with the number of NP co-expression, assuming GADs expression can represent GABA levels in the cell (111). In contrast, we found a weak but significant negative correlation between total GADs expression and NP co-expression in both controls (p<0.01; r=-0.10) and AD. (p<0.01; r=-0.19) (Fig S5). We tested the correlation separately for *GAD1* as well as *GAD2*, but a significant negative correlation only existed for *GAD1* expression (p_control_<0.0001, r=-0.14; p_AD_<0.05, r=-0.15). Our findings suggest that GABAergic neurotransmission does not intrinsically confer the selective vulnerability hypothesized for the disproportionate loss of HNP cells observed in AD brains here. In short, we discovered that there was a disproportionate absence of HNP cells in the EC of AD brains. We also found that a greater number of co-expressed NPs, but not the level of GABA, is correlated with the loss of HNP neurons in AD.

### Expressions of ADNP genes decrease with age in the hippocampus

Advanced age is the biggest risk factor for developing AD, however, even normal aging (counter to pathological aging) is accompanied by a degree of cognitive decline as the volume of gray matter and the number of neurons in the medial temporal lobe–including hippocampus and EC–decrease moderately with age (112–114). To show that the abnormal decrease in NP expression in AD EC could play a role in the early pathogenic process of AD, we examined the gene expression changes of AD-associated NPs (ADNP) identified previously in the general aging population. We hypothesized that if the decrease in ADNP expression contributed to the early pathology of AD, we would observe a decrease in ADNP expressions with aging (a negative correlation between ADNP expression and age) specifically to EC and brain regions near EC, such as the hippocampus. To test this hypothesis, we downloaded tissue-specific RNA-seq data, including those generated from the hippocampus (n=74) and pancreas (n=141), obtained by the GTEx consortium, and analyzed the age-related characteristics of ADNP expression in these tissues (41). RNA-seq data from the pancreas was included in the analysis as it is a peripheral organ outside the CNS and pancreatic cells transcribe many NP genes included in this study. We expected that the gene expressions from pancreatic tissues would not have the hypothesized age-related correlation because although evidence exists to suggest an interplay between AD and the altered metabolic profile of pancreatic tissues (115), the hallmarks of AD are neuropathological changes in the EC and hippocampus. As expected, we found a significant negative correlation (r=-0.32; p<0.01) between the total expression of the ADNPs in the EC and age only in the hippocampus but not the pancreas (r=-0.022; p=0.7) (Fig 5A). Because it is known that many gene expressions alter with age (116), the observation could reflect the intrinsic difference between tissues rather than ADNP expression profiles. To overcome this possibility, we compared the ADNP-age correlation coefficients from the targeted tissue to those formed by random combinations of gene expression with age in the same tissue type. Again, we only found more negative ADNP-age correlation coefficients (p<0.05) in the hippocampus, suggesting that the overall ADNP expression decreases with age in the hippocampus but not the pancreas (Fig 5B). Further, we obtained the correlation coefficients and tested the correlation between the expression of each ADNP gene and age. We assigned each gene a class of correlation type (significant negative) based on the correlation coefficients and their statistical information. As shown in figure 5C, ~80% of ADNP genes expressed in the hippocampus showed a negative correlation with age while ~55% is the case for the pancreas. In addition, far more ADNP gene expressions exhibited a statistically significant negative correlation with age in the hippocampus than pancreas (18 vs 2; Table S9). *PTHLH* shared the strongest negative correlation in the hippocampus with age (r=-0.40; p<0.001) and not in the pancreas (Fig 5D). Similarly, several other ADNP genes identified previously from the single-nuclei data, including *VIP* and *CCK*, were highlighted here, negatively correlating with age only in the hippocampus (r=-0.34; p<0.01; r=-0.28; p<0.05) (Fig 5D). In summary, we report that ADNPs identified previously from the AD transcriptome analysis demonstrated age-related changes specific to the hippocampus, suggesting their roles in cognitive decline and early pathogenesis of AD.

**Fig 5.**
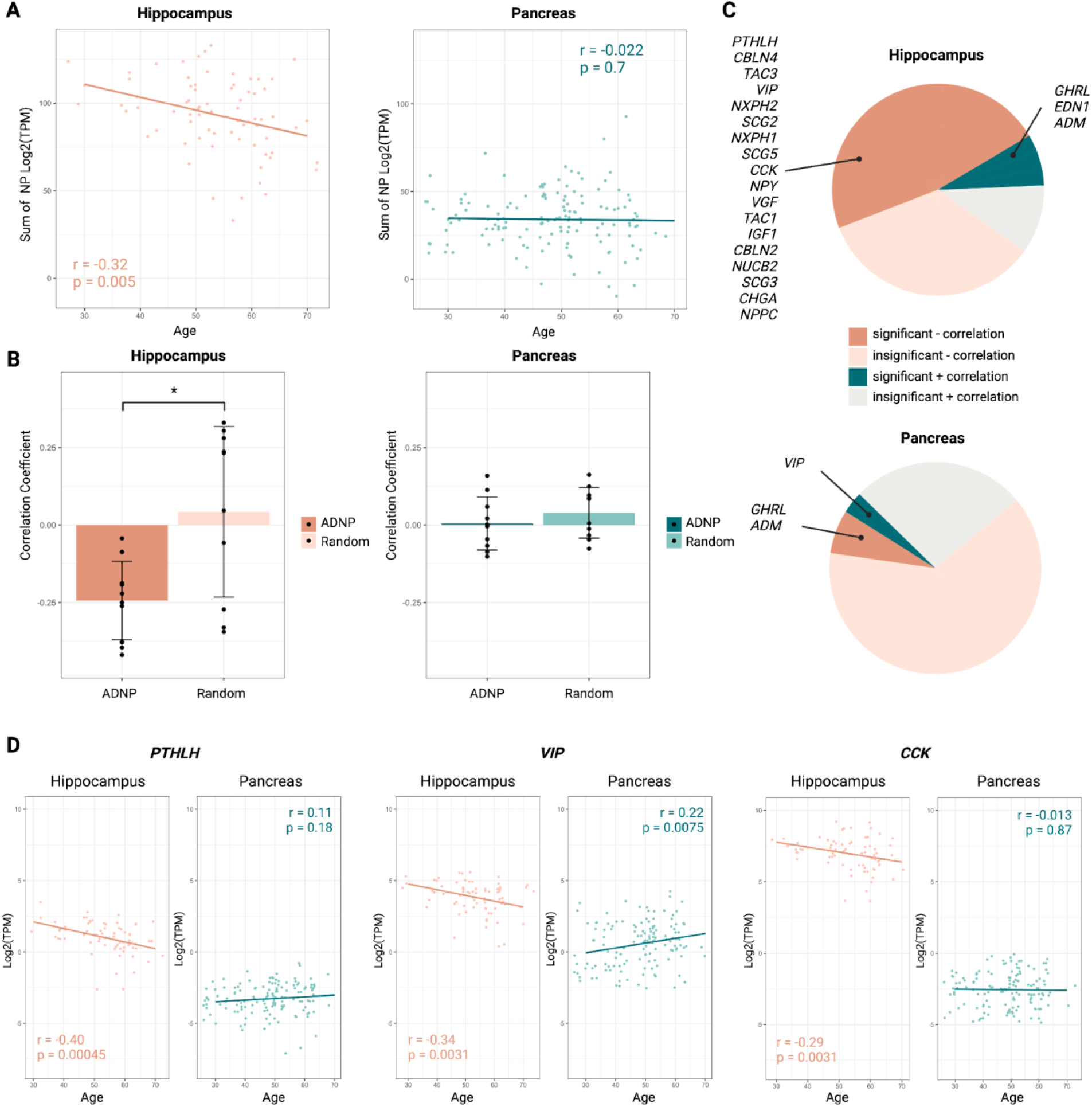
Age-related decrease of ADNP expression is unique to the hippocampus. (A) The correlation between the expression level of ADNP (sum of relative transcript count) and age was explored for the hippocampus and pancreas. (B) The correlation coefficients derived from the correlation between the expression level of random ADNP vs non-ADNP genes and age were compared in the hippocampus and pancreas. One-tailed Wilcoxon signed-rank test was used. *, p<0.05. (C) ADNPs with a significant correlation between expression level and age are highlighted for the hippocampus and pancreas. +, positive; -, negative. (D) The correlations between the expression of exemplary ADNP genes (*PTHLH, VIP, CCK*) and age are shown for the hippocampus and pancreas.

## Discussion

In the present study, we conducted a thorough analysis of the neuropeptide (NP) signaling network using the single-cell transcriptome data generated from Alzheimer’s disease (AD) patient and control brains (23), seeking to characterize cellular heterogeneity and dysfunction of the NP signaling network in the entorhinal cortex (EC) of human AD brain.

We first reported that there was a decreased expression of several neuropeptides across various cell types in AD at the single-cell level. As we were preparing this manuscript, Podvin et al. reported dysregulation of neuropeptides in the temporal cortex of AD brains using global neuropeptidomics analysis of cortical synaptosomes (117). The neuropeptides highlighted in their manuscript, including CCK, VGF, and those in the chromogranin-secretogranin family, were also identified at the transcriptomic levels in our analysis. We recognize that the data analyzed here is entirely comprised of transcriptomics, therefore we are unable to draw any definitive conclusions about NP alterations at the protein level, especially specific isoforms or their ratios of NPs that require post-translational modifications. However, we are confident that the decrease in NP expression that we observed here is reflected at the protein level, too, as many previous studies including Podvin et al. have shown such abnormalities dispersedly (19,117).

Despite limitations intrinsic to studying transcriptomics, the NP signaling network disruption characterized by our study remains significant. Firstly, previous studies generally assumed that neurons are the source of detectable NPs, therefore, did not touch upon the source of NP changes. We showed that although neurons showed the most salient decrease in NP synthesis, the reduction of NP transcription is not limited to neurons in AD. This is especially noticeable for OPCs as the decrease in the proportion of cells expressing NPs was almost comparable to neurons in AD. To complement reports of NP abnormalities in the existing literature, we presented a fuller picture of the disrupted NP signaling network by investigating both NPs and their GPCRs. Our results brought forth the remarkable notion that abnormal expression of NP likely occurs earlier than alterations of GPCRs in AD and that the changes in the expression of GPCRs are likely adaptive to and downstream of the decrease in NP production. Moreover, NP signaling is highly redundant, therefore dispersed reports of NP abnormality from different studies without in-group comparisons cannot pinpoint the key NPs that are disturbed in AD. We overcame this issue and identified a list of AD-associated NPs (ADNPs) by examining the full range of existing NPs in single-cell data analysis. In confirming our findings by applying ML to bulk RNA-seq from two independent studies with a larger sample size, we identified model-driving ADNPs that can direct future empirical studies.

We note that the observed decrease in NP expression reported here could be an underestimation of the true changes occurring in AD EC cells. In our analysis, ~80% of neurons express at least one major type of neuropeptide in control subjects; this number was hypothesized to be 100% in mice based on the cortical neuropeptide single-cell analysis conducted by Smith et al (10). This discrepancy could be attributed to many factors, such as sample preparation, sequencing protocols, and cell identification. However, we think the most probable cause was that the data analyzed by Smith et al had a better sequencing coverage and depth for neurons and a biased selection of high peptide expression subtypes as it was enriched in neurons and especially *Vip*, *Sst*, and *Ndnf* (partly *Npy*) cells due to the FACS sorting process (10). The data analyzed here generated by Grubman et al (2019), on the other hand, could reflect a more physiological proportion of cell types, allowing us to survey the NP expression more closely to physiological and pathological conditions among different cell types in control and AD brains (23).

Further, we identified a disproportionate absence of high NP-producing (HNP) cells in the EC of AD subjects. This novel observation fits well and provides additional insights into the multifaceted pathophysiology of AD. Agreeing with decades of research before, neurons are the most active in producing NP and the HNP cells are mostly GABAergic neurons. We add to the existing body of knowledge that the abundance of NP transcripts and the number of NP expressed in a single brain cell share a strong positive correlation. In a steady state, as reasoned by Smith et al, high NP transcripts in a single cell often imply a high overall rate of processing and secretion of the active NP. It was recently shown that maintaining synaptic vesicle (SV) load at synaptic terminals in hippocampal neurons comes at a high energy cost partially due to vacuolar-type ATPases (108). Since NPs are mostly stored in dense core vesicles and often co-released with other neurotransmitters in SV, it is reasonable to hypothesize that storing and maintaining NP at synaptic terminals and/or other parts of the cell would also be metabolically demanding. In this sense, our study suggests the existence of specialized groups of HNP neurons and cells in the human CNS facing high metabolic demands and are, therefore, metabolically vulnerable for synthesizing, storing, and secreting NP to maintain homeostasis in the EC, if not also other regions of the CNS. It was then not surprising to find that neurons normally co-expressing above four NPs were completely absent in all AD brains included in the single-cell analysis, implying a selective loss of cells with such metabolic vulnerability. The increased overall production of GPCRs in different cell types, also described in this analysis, could then be interpreted as an adaption of brain cells to compensate for the loss of NP signaling. Future empirical investigations focusing on whether high NPs confer metabolic vulnerability to cells in the CNS are required to confirm or refute these hypotheses.

The intertwined relationship between high NP transcription and GABA synthesis is another point of interest. Early GABAergic system dysfunction has been shown in multiple animal studies and human subjects to underlie early neural network hyperactivity and memory impairments in AD (106). While research efforts have focused on identifying GABAergic vulnerability in AD have produced valuable understandings of the GABAergic microcircuitry and synchronized neuronal activity in the hippocampus, it remains unclear why some GABAergic neurons are more susceptible to AD than others and why administration of GABA receptor agonists has very limited if not opposite effects on improving AD symptoms (106,118–120). If our hypothesis that HNP cells are selectively vulnerable to mitochondrial dysfunction and oxidative stress due to their high metabolic demands were to hold, it would suggest that the early loss of GABAergic neurons and other cells is a reflection, or by-product, of the loss of HNP cells–the majority of which are neurons by far. It would not be surprising then that simulating the effects of GABA, rather than the lost NPs, could not rescue neuronal loss or halt the progression of AD. Our study also underlines the challenges to characterize GABAergic vulnerability due to HNP as these GABAergic neurons account for only ~3% of all cells surveyed. Future research characterizing HNP cells in normal and AD brains may provide insights into the mechanisms contributing to the selective absence of HNP GABAergic cells in AD. The question also remains regarding why NP-producing cells in the hippocampus and entorhinal cortex would be the most vulnerable among all brain regions in AD. While we only investigated EC, research presenting a comprehensive and multi-omic single-cell atlas of NP-producing cells describing their density and activity in the human brain would begin to provide answers. We expect such studies to lead to the identification of the subgroup(s) of hippocampal GABAergic cells, especially neurons, that demonstrate uniquely high ADNP co-expression and abundance.

Cognitive decline in the very early stages of AD and during normal aging share some molecular and cellular changes. Hypothesizing that the decline of ADNPs identified in the single-cell transcriptomic data contributes to early pathogenesis in AD, we showed that there is a negative correlation between ADNP expressions and increased age existing in the hippocampus but not the pancreas. Because of the pleiotropy and redundancy of NP signaling, potential selective vulnerability thus early loss of such cells and/or the gain and loss of functions of ADNPs could explain a broad spectrum of AD pathophysiology from the prodromal to late stages, including mitochondrial dysfunction (56–60), GABAergic system impairment (3), circadian rhythm (52,61–67) and gut-brain connection disruptions (121–123), glial activation (53,68–72), as well as infection and immune system abnormalities (4,122). On the other hand, these processes that occur early in AD can also be caused by genetic predispositions and lifestyle choices (e.g., sleep deprivation, poor diet, and untreated chronic inflammation) that trigger and/or amplify the cascade of decreased NP production, leading to initiation and/or expedition of AD development. Regardless, we think that the decline of ADNPs in AD patients may be much faster than in normal aging populations later in life, which could make certain combinations of ADNPs suitable as AD biomarkers (Fig 6). To test such a hypothesis and promote the development of AD biomarkers, prospective longitudinal studies measuring multiple ADNP levels in various tissues assisted by machine learning are likely required.

Lastly, we note that many receptors for ADNPs are GPCRs, which have long been thought to be the most attractive targets for psychiatric drug development due to their ease of access and diversified downstream signaling pathways (11). As a result, hundreds of GPCR drugs have been approved by the FDA and many are currently in clinical trials for a wide range of conditions. However, present FDA-approved treatments for AD focus on neurotransmitters and protein misfolding processes, and strategies for NP-based treatment are yet to be developed. To demonstrate the pharmacological implications of our findings, we summarized the NP-GPCRs of differentially expressed NPs from the GPCR database (Table S10) (124). For GPCRs included by the database that are cognate receptors for ADNPs discovered here, 18 receptors are targeted by 60 different FDA-approved drugs and 9 more receptors are being targeted by 58 unique drugs currently in clinical trials (Table S10). Our study also showed some cell-type-specificity of NP expression, which could be translated to fewer side effects of NP-mimicking/GPCR-targeting drugs. When considering the increasing number of pharmaceutical agents having diverse modes of action developed for a growing number of GPCR targets, as well as the opportunity for drug repurposing, our results highlight the potential for treating AD by simultaneously targeting multiple druggable GPCRs.

To conclude, we identified ADNPs and reported their abnormal expression in AD brains by comprehensively surveying NPs and GPCRs using transcriptomic data. We revealed a disproportionate absence of cells that abundantly express multiple NPs in AD brains and proposed that there is a selective vulnerability of the subpopulation(s) of GABAergic cells that express a high quantity of NPs. We also provided evidence supporting the role of ADNPs in the early development of AD. Together, our study provides important insights that can be leveraged to assist the development of AD biomarkers and therapeutic strategies.

## Acknowledgments

Dr. Ling Li from the Department of Experimental and Clinical Pharmacology and Dr. Alice Larson from the Department of Veterinary and Biomedical Sciences at the University of Minnesota provided helpful comments that improved the quality of the research presented herein.

The results published here are in part based on data obtained from the AD Knowledge Portal (**https://adknowledgeportal.org/**). These data were generated from postmortem brain tissue collected through the Mount Sinai VA Medical Center Brain Bank and were provided by Dr. Eric Schadt from Mount Sinai School of Medicine. The Mayo RNAseq study data was led by Dr. Nilüfer Ertekin-Taner, Mayo Clinic, Jacksonville, FL as part of the multi-PI U01 AG046139 (MPIs Golde, Ertekin-Taner, Younkin, Price). Samples were provided from the following sources: The Mayo Clinic Brain Bank. Data collection was supported through funding by NIA grants P50 AG016574, R01 AG032990, U01 AG046139, R01 AG018023, U01 AG006576, U01 AG006786, R01 AG025711, R01 AG017216, R01 AG003949, NINDS grant R01 NS080820, CurePSP Foundation, and support from Mayo Foundation. Study data includes samples collected through the Sun Health Research Institute Brain and Body Donation Program of Sun City, Arizona. The Brain and Body Donation Program is supported by the National Institute of Neurological Disorders and Stroke (U24 NS072026 National Brain and Tissue Resource for Parkinson’s Disease and Related Disorders), the National Institute on Aging (P30 AG19610 Arizona Alzheimers Disease Core Center), the Arizona Department of Health Services (contract 211002, Arizona Alzheimers Research Center), the Arizona Biomedical Research Commission (contracts 4001, 0011, 05-901 and 1001 to the Arizona Parkinson’s Disease Consortium) and the Michael J. Fox Foundation for Parkinson’s Research.

The Genotype-Tissue Expression (GTEx) Project was supported by the Common Fund of the Office of the Director of the National Institutes of Health, and by NCI, NHGRI, NHLBI, NIDA, NIMH, and NINDS. The data used for the analyses described in this manuscript were obtained from the GTEx Portal (https://www.gtexportal.org/) on 10/05/2022.

Figures were organized in BioRender (biorender.com).

