## Supplementary material for "Single-cell sequencing of Entorhinal Cortex Reveals Wide-Spread Disruption of Neuropeptide Networks in Alzheimer’s Disease": SupplementaryMaterials.docx

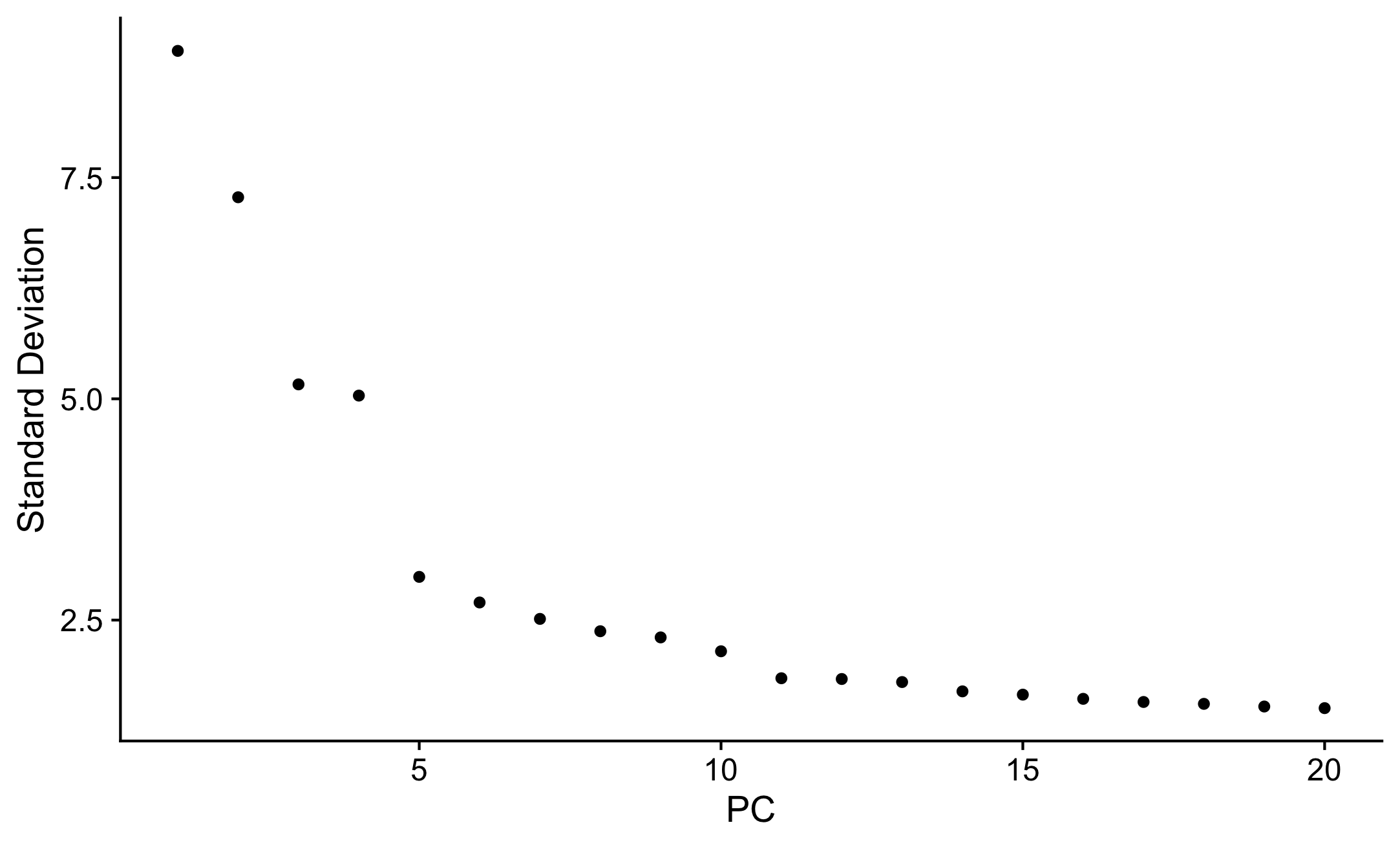

**Fig S1. Elbow plot visualizing the standard deviation of each principal component.**

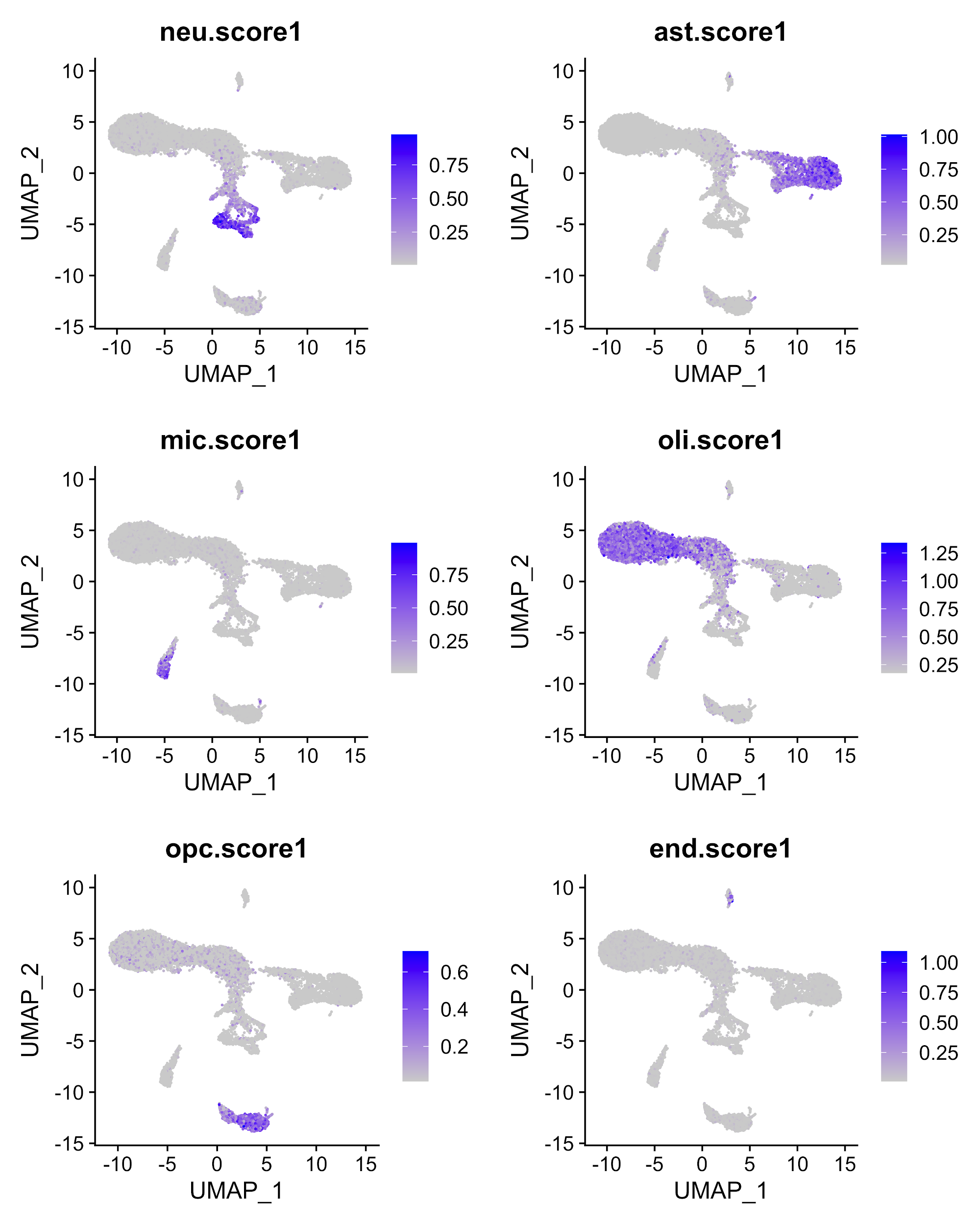

**Fig S2. BRETIGEA scoring cell clusters.** Neu.score1, BRETIGEA score for neurons; ast.score1, BRETIGEA score for astrocytes; mic.score1, BRETIGEA score for microglia; oli.score1, BRETIGEA score for oligodendrocytes; opc.score1, BRETIGEA score for oligodendrocyte progenitor cells; end.score1, BRETIGEA score for endothelial cells.

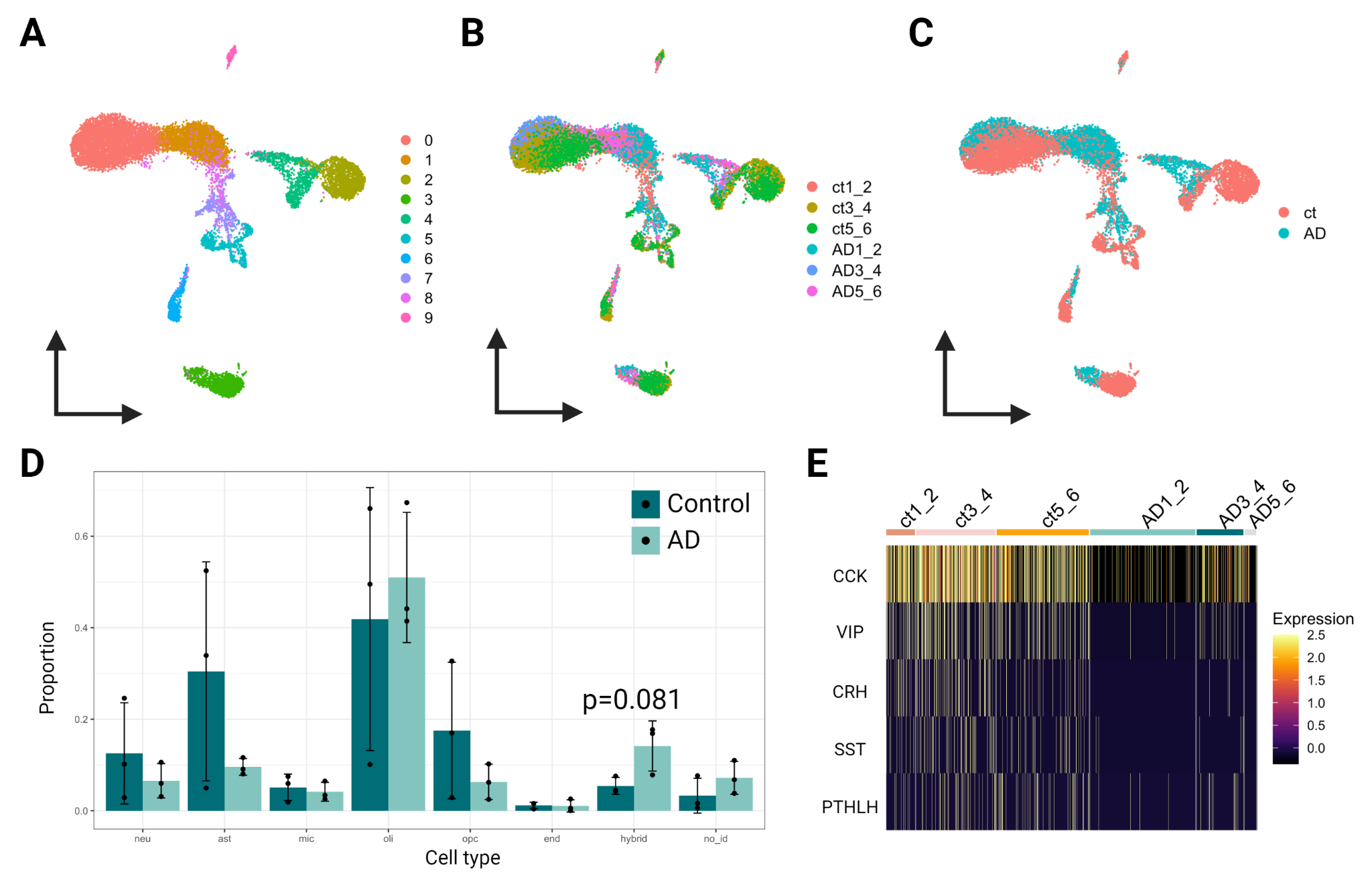

**Fig S3.** (A) Classification by the K-nearest neighbors algorithm. (B) Classification by grouped replicates. (C) Classification by phenotype. Ct, control. (D) A cell type-specific comparison of the cell composition between control and AD entorhinal cortex. Two-tailed Wilcoxon signed-rank test was used. Relative proportion = number of cells expressing gene target for each cell type per biological replicate/number of cells in the cell type per biological replicate. (E) Expression of differentially expressed NPs in neurons using the gene list by Smith et al.

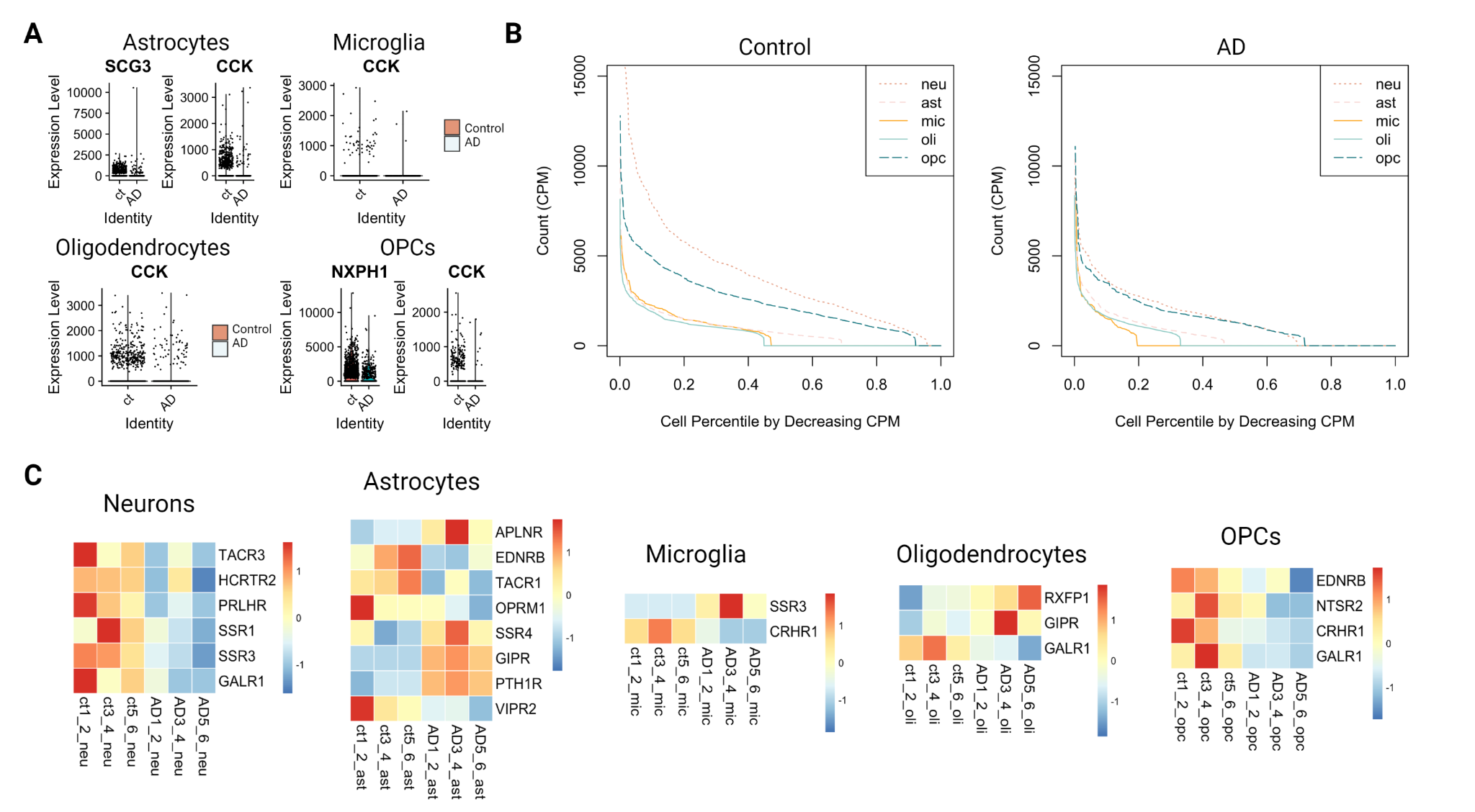

**Fig S4.** (A) Differentially expressed neuropeptide (NP) genes in astrocytes, microglia, oligodendrocytes, and oligodendrocyte progenitor cells (OPCs). Ct, control. (B) The total relative count of all selected NPs plotted against the cell percentile ranked by decreasing cell count in selected cell types in control and AD. (C) Cell type-specific heatmaps showing the G protein-coupled receptor genes expressed by significantly changed (One-tailed Wilcoxon signed-rank test, p<0.1) proportions of cells for each grouped biological replicate. Genes expressed only in one grouped replicate were excluded. Relative proportion = number of cells expressing gene target for each cell type per biological replicate/number of cells in the cell type per biological replicate.

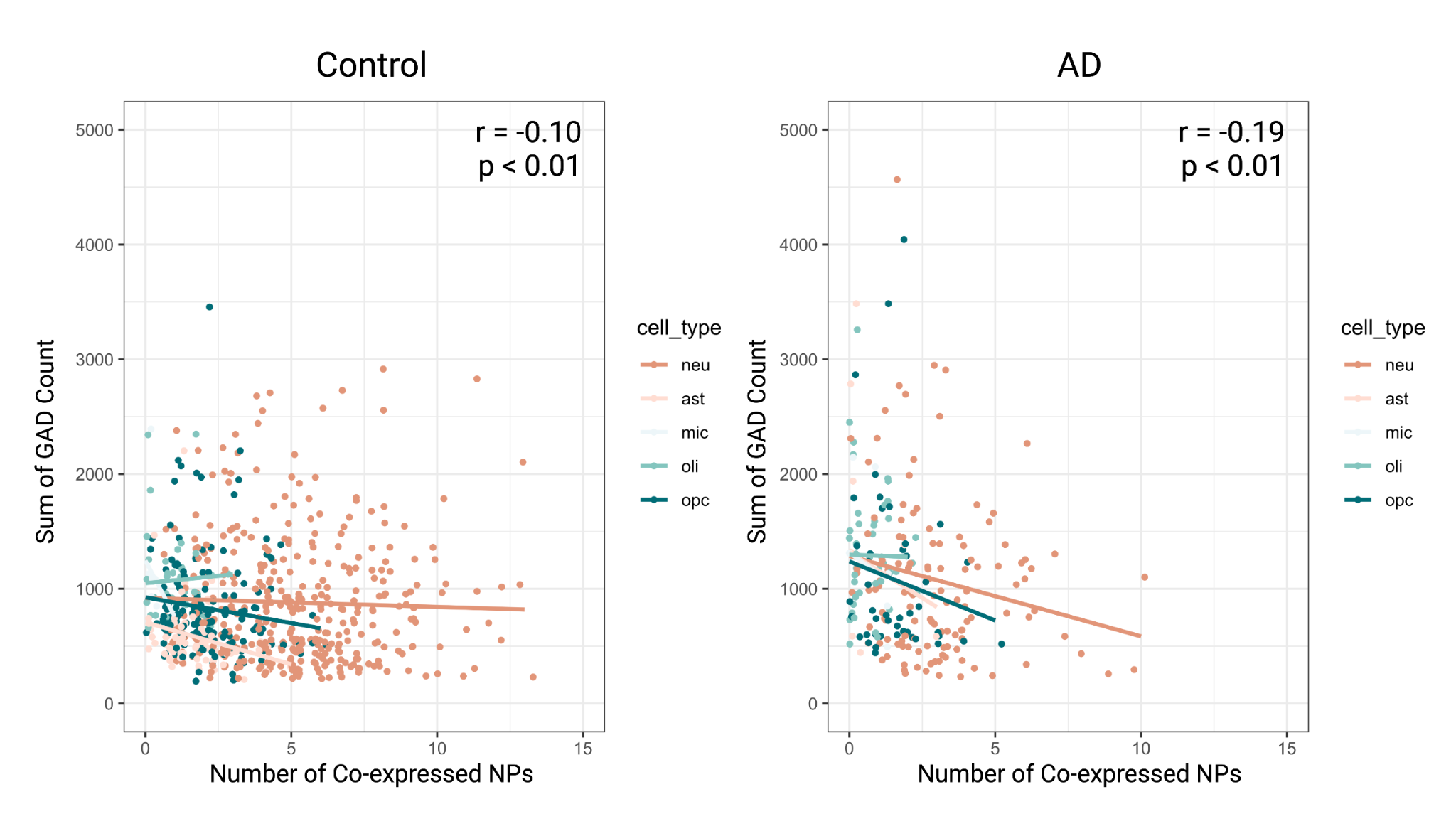

**Fig S5.** **Significant correlation between the sum of the relative number transcripts from GAD (glutamate dehydrogenase) genes and the number of coexpressed NPs shown in both control and AD EC cells.** Pearson's test of correlation was used. r, correlation coefficient.
